# Transcriptional profiling of mucus production and modification in rhesus macaque endocervical cells under hormonal regulation

**DOI:** 10.1101/2023.05.18.541362

**Authors:** Katrina Rapp, Shuhao Wei, Mackenzie Roberts, Shan Yao, Suzanne S. Fei, Lina Gao, Karina Ray, Alexander Wang, Rachelle Godiah, Leo Han

**Author notes:** **Corresponding author:** Leo Han MD MPH, Oregon National Primate Research Center, 505 NW 185^th^ Ave, Rm 2116, Beaverton, OR 97006. **Disclosure statement:** Katrina Rapp: None, Shuhao Wei: None, Mackenzie Roberts: None, Shan Yao: None, Suzanne Fei: None, Lina Gao: None, Karina Ray: None, Alexander Wang: None, Rachelle Godiah: None, Leo Han: None. **Attestation statement:** - Data regarding any of the subjects in the study has not been previously published unless specified. - Data will be made available to the editors of the journal for review or query upon request. **Capsule:** Using primary epithelial endocervical cells, we profile *in vitro* hormonal regulation of mucus production, modification, and secretion.

## Abstract

**Objective:** Endocervical mucus production is a key regulator of fertility throughout the menstrual cycle. With cycle-dependent variability in mucus quality and quantity, cervical mucus can either facilitate or block sperm ascension into the upper female reproductive tract. This study seeks to identify genes involved in the hormonal regulation of mucus production, modification, and regulation through profiling the transcriptome of endocervical cells from the non-human primate, the Rhesus Macaque (Macaca mulatta).

**Design:** Experimental.

**Setting:** Translational science laboratory.

**Intervention:** We treated differentiated primary endocervical cultures with estradiol (E2) and progesterone (P4) to mimic peri-ovulatory and luteal-phase hormonal changes. Using RNA-sequencing, we identified differential expression of gene pathways and mucus producing and modifying genes in cells treated with E2 compared to hormone-free conditions and E2 compared to E2-primed cells treated with P4.

**Main Outcome Measures:** We pursued differential gene expression analysis on RNA-sequenced cells. Sequence validation was done using qPCR.

**Results:** Our study identified 158 genes that show significant differential expression in E2-only conditions compared to hormone-free control, and 250 genes that show significant differential expression in P4-treated conditions compared to E2-only conditions. From this list, we found hormone-induced changes in transcriptional profiles for genes across several classes of mucus production, including ion channels and enzymes involved in post-translational mucin modification that have not previously been described as hormonally regulated.

**Conclusion:** Our study is the first to use an *in vitro* culture system to create an epithelial-cell specific transcriptome of the endocervix. As a result, our study identifies new genes and pathways that are altered by sex-steroids in cervical mucus production.

## INTRODUCTION

Endocervical mucus regulates fertility via changes in mucus quantity and quality throughout the menstrual cycle, allowing sperm entry only during the fertile window when ovulation occurs.^1^ During the peri-ovulatory period, high serum estradiol (E2) levels drive increased mucus production and reduced mucus viscosity, facilitating sperm entry through the cervix. Following ovulation, the luteal rise in progesterone (P4) results in increased mucus viscosity that prevents sperm and ascending pathogens from entering the upper reproductive tract.^1^ These cyclical changes in mucus have long been used by women to determine the fertile window and serve as a fundamental mechanism of action of progestin-containing contraceptives.^2^

Mucus in the female endocervix is produced by a single layer of columnar epithelium lining the endocervix, and is made of water, amino acids, lipids, enzymes, bactericidal proteins, plasma proteins, and mucins.^3^ Mucins are large glycol proteins that serve as the primary structural protein in mucus and bind water to form an intermolecular gel, with mucin 5B (MUC5B) acting as the key gel-forming mucin within the endocervix.^3, 4^ Ion channels have also been identified as critical determinants of mucus characteristics, as they produce an ionic milieu that facilitates water entry into the lumen and the unfolding of mucin oligomers.^5^ Ion channels such as the cystic fibrosis transmembrane conductance regulator (CFTR) anion channel, anoctamin-1 calcium-activated chloride channel (ANO1), and sodium epithelial ion channel (ENaC) have all been demonstrated to lead to abnormal mucus production when inhibited or knocked out.^6-8^ In addition to the luminal hydration of mucus, other identified components of mucus physiology include proteins that mediate post-translational modification, packaging and release of mucins, and enzymes that digest mucin oligomers.^9, 10^

To understand how hormone changes throughout the menstrual cycle regulate cervical mucus characteristics, we developed an *in vitro* primary endocervical cell culture model using conditionally reprogrammed cells (CRCs) from the non-human primate model of Rhesus Macaques (Macaca mulatta). Macaques possess homologous structure and function of the cervix to humans, including a mucus-secreting endocervix. Only primates have menstrual cycles, making them an ideal animal model for studies related to endocervical physiology. By using an *in vitro* model, our transcriptome focuses on the epithelial cells that produce mucus, as the majority of cells in the endocervix are non-epithelial.^11^ We previously established that CRC methodology enables us to overcome the limited expansion potential of endocervical cells and avoid the loss of *in vivo* characteristics, in particular hormonal sensitivity, that occurs with immortalized cell lines. The CRCs utilized in this experiment have been shown to expand and maintain steroid receptor expression and secrete mucus analogous to *in vivo* mucus, and therefore provide an ideal model for mechanistic studies of cervical mucus regulation.^12, 13^

To gain a better understanding of the molecular basis of mucus changes throughout the menstrual cycle, we performed RNA-sequencing (RNA-seq) on our cultured endocervical cells. From that data set, our study specifically explores the hormonal regulation of pathways and genes involved in mucin synthesis, mucus hydration via ion channel regulation, mucus cross-linking and stabilization, protein- and enzyme-mediated mucus modification, motile ciliogenesis, and hormonal signaling (Figure 1).

**Figure 1.**
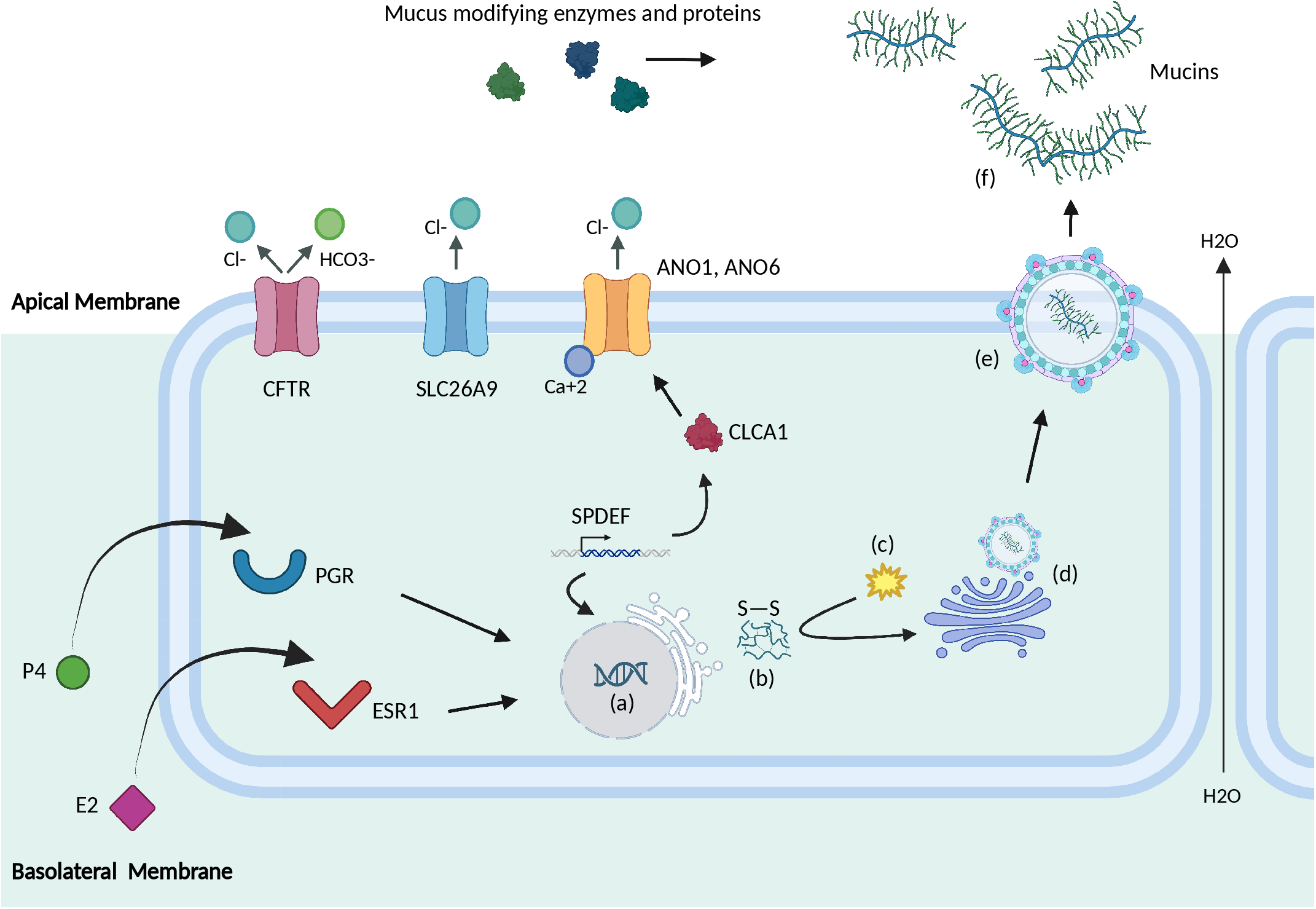
Mucin biosynthesis is hormonally regulated by E2 and P4. E2 and P4 levels regulate transcription of mucin genes and genes that influence mucin character and structure (a). Mucin peptides undergo folding and disulfide bond formation (b) followed by glycosylation (c). Mucins are then packaged in the Golgi apparatus (d) and exported into the lumen via SNARE-mediated membrane fusion (e). Once outside the cell, mucin proteins are further modified by enzymes and proteins and expand in the presence of water, which is increased in the presence of luminal anions such as Cl^-^ and HCO_3_^-^ (f). Created with BioRender.com

## METHODS

### Conditionally Reprogrammed Endocervical Cells

The Oregon National Primate Research Center (ONPRC) provided care and husbandry for the Rhesus Macaques in this study. All animal study procedures were reviewed and approved by the ONPRC Institutional Animal Care and Use Committee (IACUC). Conditional reprogramming to produce primary endocervical cell cultures (CRECs) from reproductive-aged female Rhesus Macaques (Macaca mulatta) has been previously described in detail.^12^ Briefly, cervical tissues from animals undergoing necropsy for studies unrelated to our own were obtained. Endocervical epithelial tissue was isolated, digested, and co-cultured in conditionally reprogrammed cell conditions to propagate and expand the primary endocervical cells.^12, 14^ To differentiate cultured cells, we seeded 1 × 10^5^ cells on permeable supports (12mm diameter, 0.4μM pore size, Costar Transwell, Corning, Action, MA, USA) in calcium (Ca^+2^) supplemented media (total concentration = 0.4 mM, Sigma Aldrich, St. Louis, MO, USA).

### Hormone Treatments

Differentiated primary endocervical cells were treated with hormonal conditions that mimic periovulatory and mid-luteal conditions *in vitro*.^12, 14, 15^ For peri-ovulatory conditions (abbreviated as EE), E2 was added to differentiation media once daily for nine days (10−8 M, Sigma Aldrich). For luteal conditions (abbreviated as EP), cells were initially primed with E2 (10−8 M) for seven days, followed by P4 (10−7 M, Sigma Aldrich) and E2 (10−9 M) for 48 hours. We chose a relatively short P4 treatment as clinical studies suggest mucus changes occur rapidly with progestin administration.^16, 17^ We also had a vehicle-only control that served as our hormone-free comparator (abbreviated as CC). Endocervical cell cultures used in RNA-seq originated from four distinct female animals, ages 4 to 16 years old, with three to five separate experiments for each animal generated from cells differentiated between passages three and five. We performed each experiment in quadruplicate and pooled samples from similar conditions for analysis.

### RNA Isolation and cDNA Synthesis

Following hormone treatment, total RNA was extracted from cell cultures using the TRIzol RNA+ (Invitrogen, Gaithersburg, MD, USA) mini-kit following manufacturer instruction. RNA was quantified using the Nanodrop system (Thermofisher). Superscript III First Strand cDNA Synthesis kit (Invitrogen, Waltham, MA, USA) was used to generate cDNA from 1μg of RNA following manufacturer instruction.

### RNA-Sequencing

RNA was evaluated for integrity using a 2100 Bioanalyzer (Agilent, Santa Clara, CA, USA). Libraries for RNA-seq were then prepared using the TruSeq Stranded mRNA Library Prep Kit (Illumina, San Diego, CA, USA). Briefly, poly(A)+ RNA was isolated from the total RNA using magnetic beads coated with oligo(dT). The poly(A)+ RNA was then fragmented by incubation with divalent cations and heat, then converted to double stranded cDNA using random hexamer primers. The second strand was synthesized with dUTP to fix the stranded orientation of the libraries. After ligation of adapters, the library was amplified by limited rounds of polymerase chain reaction. Libraries were profiled on a 4200 TapeStation (Agilent) D1000 tape and quantified using an NGS library quantification kit (Roche/Kapa Biosystems) on a StepOnePlus real time PCR workstation (Thermo/ABI). Libraries were sequenced on a NovaSeq 6000 (Illumina). Fastq files were assembled using bcl2fastq (Illumina).

### Transcriptome Analysis

Differential expression analysis was performed by the ONPRC Bioinformatics & Biostatistics Core. The quality of the raw sequencing files was evaluated using FastQC^18^ combined with MultiQC.^19^ Trimmomatic was used to remove remaining Illumina adapters.^20^ Reads were aligned to Ensembl’s mmul10 along with its corresponding annotation, release 106. The program STAR (v2.7.3a), shown to out-perform other RNA-sequencing aligners, was used to align the reads to the genome.^21,22^ Utilizing the gene annotation file, STAR calculated the number of reads aligned to each gene. RNA-SeQC^23^ and another round of MultiQC were utilized to ensure alignments were of sufficient quality. Gene-level raw counts were filtered to remove genes with extremely low counts in many samples following published guidelines,^24^ normalized using the trimmed mean of M-values method,^25^ and transformed to log-counts per million with associated observational precision weights using voom method.^26^ Gene-wise linear models with primary variable treatment group were employed for differential expression analyses using limma with empirical Bayes moderation^27^ and false discovery rate (FDR) adjustment.^28^ To facilitate identification of functional biological modules in response to these hormonal changes, we used an FDR p-value cut-off of <0.2 for EE versus CC and an unadjusted p-value cut-off of <0.05 for EP versus EE.

### Gene Expression Analysis

To confirm expression data found in RNA-seq analysis, we measured gene expression of hormonally treated endocervical cells from additional experiments with the same hormonal treatment conditions using real-time quantitative PCR (QuantStudio 12K Flex System, Life Technologies). Mean differences in relative gene expression using ANOVA were assessed with relative gene expression as the outcome and treatment as the predictor with E2 as the referent (Figure 2). All analyses were conducted in Stata version 15.1 (Stata Corp, College Station, TX, USA).

**Figure 2.**
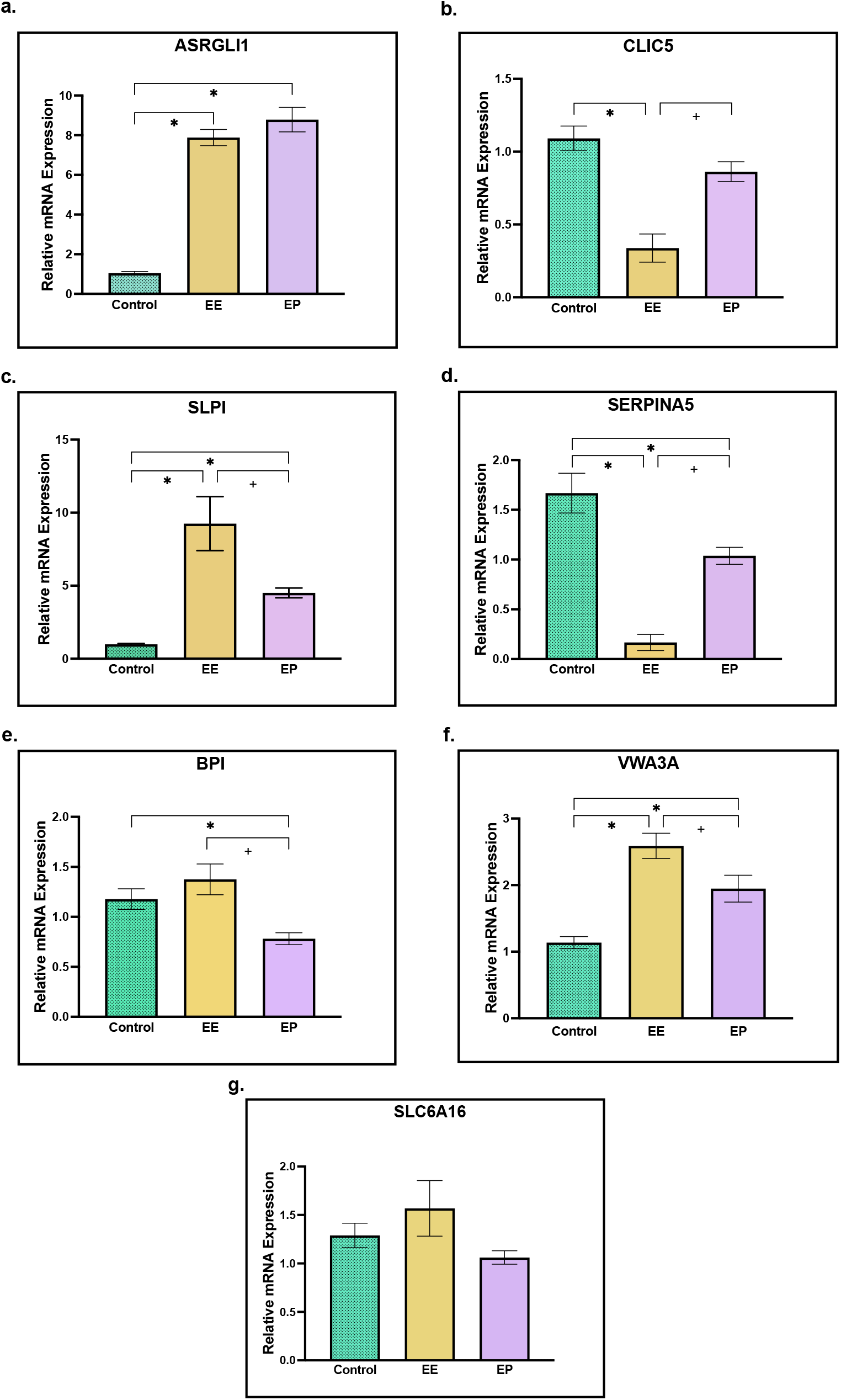
qPCR validation of RNA-sequencing results. Relative gene expression of highly up-regulated or down-regulated genes in EE vs CC or EP vs EE, including a) ASRGLI1, b) CLIC5, c) SLPI, d) SERPINA5, e) BPI, f) VWA3A g) SLC6A16. (ANOVA, p<0.05).

## RESULTS

### RNA-Sequencing Data Analysis

13,784 genes from Rhesus Macaque endocervical cell cultures were included in differential expression analysis after low count filtering. Our study identified 158 genes that show significant differential expression in E2-only conditions compared to hormone-free control, and 250 genes that show significant differential expression in P4-treated conditions compared to E2-only conditions. We then produced a list of genes and gene classes associated with epithelial mucus production and secretion throughout the menstrual cycle (Table 1).^7-10, 29-31^ From this list we found 28 differentially expressed genes (DEGs) in estradiol-only conditions (EE vs CC) and 10 DEGs in progesterone-treated conditions (EP vs EE) (Figure 3), with multidimensional scaling illustrating alterations in gene expression among our samples and conditions (Figure 4). These genes are subdivided into classes based on their role in mucus synthesis and modification, consisting of mucins, ion channels, post-secretory mucus modifiers, mucus modifying enzymes and proteins, cilia-associated genes, cell signaling genes, and aquaporins.

**Table 1.**
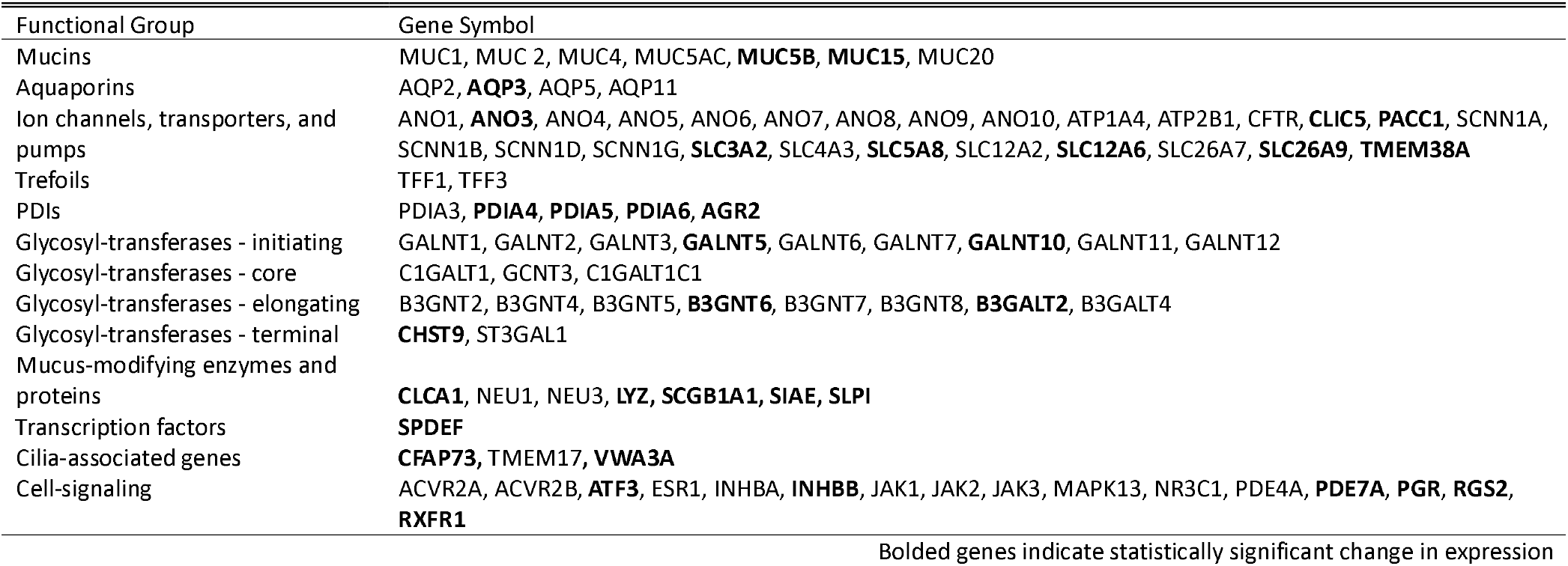
List of genes involved in mucus synthesis and modification identified in Rhesus Macaques endocervical cells

**Figure 3.**
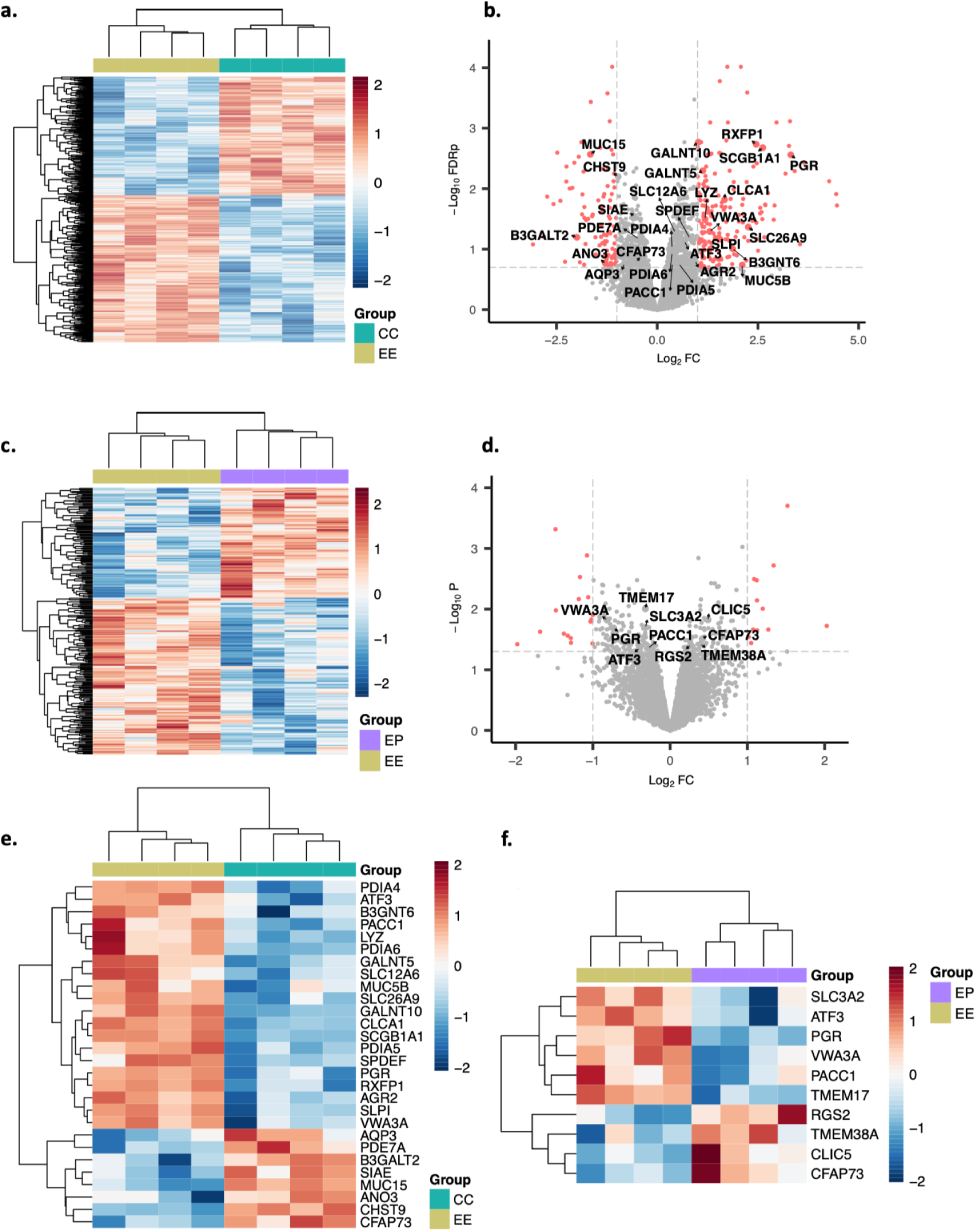
Gene expression profiling of the endocervical cells in E2-only conditions versus hormone-free control (EE vs CC) and P4-treated cells vs E2-only conditions (EP vs EE). **a**) Heat map representation of relative expression levels of genes in the endocervix samples in EE vs CC. Hierarchical cluster analysis of top differentially expressed genes was performed using the pheatmap package (RRID: SCR_016418). Each horizonal line represents a single gene and each column represents a single sample. The relative expression of each gene is color-coded as upregulated (red) to downregulated (blue). **b**) Volcano plot representation of Log_2_ fold change in expression levels in the endocervix samples in EE vs CC, highlighting genes implicated in mucus synthesis and modification. **c**) Heat map representation of relative expression levels of genes in the endocervix samples in EP vs EE. **d**) Volcano plot representation of Log_2_ fold change in expression levels in the endocervix samples in EP vs EE, highlighting genes implicated in mucus synthesis and modification. **e**) Heat map representation of relative expression levels of genes involved in mucus synthesis and modification in EE vs CC. **f**) Heat map representation of relative expression levels of genes involved in mucus synthesis and modification in EP vs EE.

**Figure 4.**
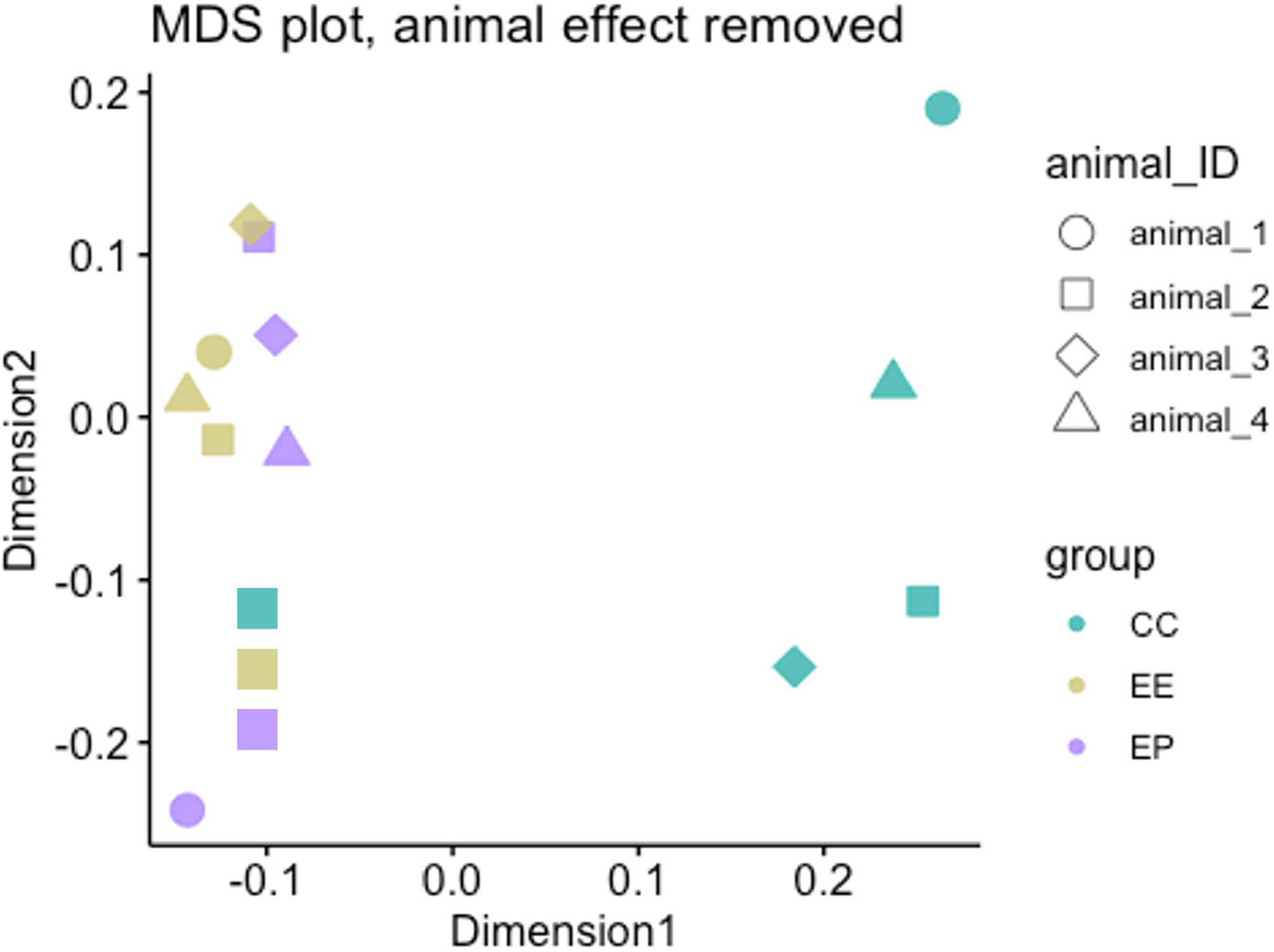
Multidimensional scaling plot showing variation in gene expression among samples and treatment group, including hormone-free controls (CC), E2-treated cells (EE) and E2-primed P4-treated cells (EP).

### Pathway Analysis

Overarching cellular pathways affected by hormonal treatment were captured through functional annotation clustering analysis using DAVID^32, 33^ and Reactome^34^ software tools. Significant pathways in E2-only conditions include the estrogen signaling pathway, ion channel transport, post-translational protein phosphorylation, glycosaminoglycan, and mTOR signaling pathway. Significant pathways in both E2-only and P4-treated conditions include metabolic pathways, WNT ligand biogenesis and trafficking, Nuclear Receptor transcription pathway, PI3K events in ERBB4 signaling, and RAF/MAP kinase cascade.

### Mucins

Significant overlap exists between mucin genes expressed in our results and those previously described (Table 1).^35^ MUC5B was the dominant mucin gene transcribed in our cells and was upregulated over 4-fold in E2-only conditions compared to hormone-free control. MUC15, a cell surface mucin that serves to create a protective mucus layer in various tissue types,^36, 37^ was significantly downregulated in E2-only conditions.

### Ion Channels, Transporters, and Pumps

Ion channels implicated in hormonally regulated mucus hydration in E2-only conditions include solute carrier family members (SLC26A9, SLC12A6) and proton-activated chloride channel 1 (PACC1). Most notably, anion transporter SLC26A9 was upregulated almost 5-fold in E2-only conditions. PACC1 was upregulated in E2-only conditions and downregulated with P4-treatment. Genes that were downregulated during E2-driven mucus hydration or upregulated during P4-driven mucus thickening include an additional member of the solute carrier family (SLC3A2), a member of the anoctamin family (ANO3), cation channel TMEM38A, and intracellular Cl^-^ channel CLIC5 (Figure 3).

### Post-Secretory Mucus Modifiers

Families of genes associated with post-secretory mucus modification and stabilization such as the protein disulfide isomerase (PDI) family (PDIA4, PDIA5, PDIA6, and AGR2) were upregulated in E2-only conditions. Additionally, several genes encoding glycosyltransferases (GALNT5, GALNT10, and B3GNT6) required for the glycosylation of mucins were upregulated in E2-only conditions (Figure 3).

### Mucus Modifying Enzymes and Proteins

Genes responsible for mucus-modifying enzymes and proteins, such as secretory leukocyte protease inhibitor (SLPI), chloride channel accessory 1 (CLCA1), secretoglobin family 1A member 1 (SCGB1A1), lysozyme (LYZ), and sialate O-acetylesterase (SIAE) showed hormonally driven alterations in expression (Figure 3). SLPI is abundant in cervical mucus during ovulation and is believed to protect sperm from the effects of elastase and therefore increase sperm penetrability through human cervical mucus.^38, 39^ SLPI was upregulated almost 3-fold in E2-only conditions. CLCA1, which encodes an accessory protein that supports ion channel conductance involved in mucus hydration,^40^ was upregulated over 3-fold in E2-only conditions. SCGB1A1, a gene implicated in attenuation of lipopolysaccharide-stimulated MUC5AC production in human bronchial epithelial cells,^41^ was upregulated over 6-fold in E2-only conditions. We also found upregulation of antibacterial enzyme LYZ and downregulation of SIAE, a modifying enzyme of sialic acid, in E2-only conditions.

### Cilia-Associated Genes

Cilia aid the movement of mucus, which is directly influenced by the hydration status of mucus.^42^ Two genes that regulate motile ciliogenesis, Cilia and Flagella Associated Protein 73 (CFAP73) and Von Willebrand Factor A Domain Containing 3A (VWA3A) showed inversed expression in E2-only and P4-treated comparisons (Figure 3).

### Cell Signaling Genes

Analysis of genes involved in cell signaling support that our EE and EP conditions are reflective of the peri-ovulatory and luteal phases of the menstrual cycle respectively. The progesterone receptor (PGR) was upregulated almost 10-fold in E2-only conditions and downregulated with P4-treatment, reflecting what is seen physiologically in the reproductive cycle.^43^ Inhibin-B (INHBB) was upregulated 2-fold in E2-only conditions, again reflecting what is seen physiologically prior to ovulation. Further, we found hormonal regulation of genes that have not been identified to play a role in fertility at the level of the cervix, despite having elucidated roles in other reproductive organs including the ovary and uterus. These include relaxin family peptide receptor 1 (RXFP1), activating transcription factor 3 (ATF3), and regulator of G-protein signaling 2 (RGS2).

### Aquaporins

Several aquaporins were expressed in our endocervical cells (Table 1). AQP3, implicated in CFTR activated water efflux in Chinese hamster ovary (CHO) cells,^44^ showed significant downregulation in E2-only conditions.

### Real-Time PCR Validation of RNA-Sequencing Results

Using real-time qPCR, we performed validation experiments of genes in our sequencing data set including SLPI, ASRGLI1, SERPINA5, SLC6A16, CLIC5, VWA3A, BPI and SLC6A16 (Supplemental Figure 1). Gene profile of tested genes match the RNA-seq profile.

## DISCUSSION

In this transcriptome study of primary endocervical epithelial cells, genes associated with mucus production showed altered gene expression in response to hormonal changes that correlate to the high estradiol levels found midcycle and the effects of progesterone in the early luteal phase. We found hormonal regulation of the gel-forming mucin MUC5B, various ion channels influencing mucus hydration, and genes supporting mucus formation. Genes involved in ovarian cell signaling during the peri-ovulatory period were also found to be under hormonal regulation in the endocervix, a new finding that warrants further investigation.

Our study emphasizes the importance of mucin-ion interactions, aligning with previous mucus research. The interaction between mucin proteins, water, and the ionic environment of the lumen is a crucial determinant of mucus properties.^5, 45, 46^ While there are conflicting reports regarding whether mucin proteins themselves are hormonally regulated,^47, 48^ our study found upregulation of MUC5B in E2-only conditions, suggesting increased mucin production influences mucus changes during this period. This is consistent with the grossly observed phenomena that mucus quantity is greatly increased during the peri-ovulatory period, where decreased viscosity suggests a higher ratio of water relative to mucin protein.^1, 49^

We identified hormonal regulation of ion channels involved in chloride (Cl^-^) transport, including SLC26A9, a promising therapeutic target for cystic fibrosis.^50^ A defective CFTR channel in the setting of cystic fibrosis results in altered Cl^-^ and bicarbonate (HCO_3_^-^) secretion, creating inadequately hydrated, viscous mucus that obstructs secreting organs.^51^ SLC26A9 functions as an alternative Cl^-^ channel and prevents mucus plugging within the airways in the setting of mucus hypersecretion.^6, 50, 52^ Therefore, upregulation of SLC26A9 in E2-only conditions favors mucus hydration and the thinned mucus observed clinically midcycle. Calcium activated Cl^-^ channels (CaCCs) have increasingly defined importance in mucosal systems, and members of the anoctamin family of CaCCs contribute to abnormal mucus production when knocked down.^53^ Interestingly, downregulation of ANO3 in E2-only conditions suggests an alternative role for this member of the anoctamin family in mucus hydration. Given the ability of ANO3 to regulate various other ion channels via scramblase activity,^54-56^ it may downregulate endogenously expressed ion channels involved in mucus hydration, therefore requiring downregulation during ovulation. Our study also found downregulation of SLC3A2 with E2-only treatment, despite its proposed supportive role to CaCCs that facilitate water transport.^57,58^ Furthermore, we identified new hormonally controlled ion channels that function to transport Cl^-^ or cations, which contribute to the ionic environment impacting mucus hydration. TMEM38A, CLIC5, PACC1, SLC3A2, and SLC12A6 exhibited variable patterns of expression in E2-only and P4-treated conditions, and one possible explanation is that fertile mucus production during the peri-ovulatory period requires downregulation of some ion channels and upregulation of others to optimize the luminal ionic content during the fertile window.

Many proteins and enzymes affect mucus hydration by altering ion channel activity and directly interacting with mucins. CLCA1 is implicated in excess mucus production when overexpressed and induces gel-forming mucin MUC5AC expression in the endocervix.^40, 59, 60^ We found upregulation of CLCA1 in E2-only conditions, supporting its role in cervical mucus production. SPDEF, a transcription factor that induces expression of CLCA1, MUC5B, and MUCAC,^61^ was also upregulated in E2-only conditions. Our results also highlight a role for SCGB1A1 in cervical mucus production. SCGB1A1 expression is implicated in bronchial epithelial MUC5AC production, and furthermore, under-expression is implicated in inflammatory airway diseases.^41, 62, 63^ Upregulation of SCGB1A1 in E2-only conditions suggests a possible supportive role to mucin production and mucus hydration in endocervical cells.

Protein disulfide isomerases (PDIs) are essential to the oligomerization of mucins and facilitate folding and chaperoning of proteins within the endoplasmic reticulum (ER).^64^ Secreted mucins are assembled into homo-oligomers via intermolecular disulfide bond formation facilitated by PDIs, and form a linear assembly of mucin subunits.^65, 66^ Increased expression of these genes (PDIA4, PDIA5, PDIA6, and AGR2) in E2-only conditions is consistent with prior findings and support the role of PDIs in mucus formation.^10^ Secreted mucins are also N-glycosylated by glycosyltransferases in the ER, which stabilize their confirmation and create negatively charged branches for binding water. Alterations in glycosylation at ovulation may promote sperm penetration,^9^ and subsets of glycosyltransferase groups showed variable expression patterns in our sequencing results. More work is needed to understand how hormonal regulation influences glycosyltransferase function in mucus hydration and viscosity.

Several genes were found to be hormonally regulated despite their unclear role in the endocervix. Relaxin, produced by the corpus luteum, interacts with RXFP1 to promote follicle growth, ovulation, and embryo implantation.^67^ While relaxin activity in the cervix has not previously been defined, RXFP1 was upregulated with E2-only treatment, indicating a possible role for relaxin and RXFP1 in fertility beyond the uterus. ATF3, which plays a role in luteal regression, was upregulated during E2-only conditions and downregulated during P4-treated conditions, likely due to the requirement of P4 to maintain the corpus luteum.^68^ As P4 levels fall in the absence of beta-HCG from an implanted embryo, ATF3 can be transcribed to facilitate regression of the corpus luteum.^68^ RGS2 is involved in signaling pathways essential to ovulation,^69^ and increases in the ovary following an ovulatory dose of human chorionic gonadotropin or luteinizing hormone.^43^ RGS2 was one of the few genes that we found significantly increased in P4-treated conditions. Interestingly, neither RGS2 nor ATF3 have defined functions in the cervix. PDE7A, which catalyzes cyclic nucleotides, was downregulated in E2-only conditions. While its role in the cervix is unclear, we know that PDE family members activate ion channels, such as in the case of PDE4 and CFTR.^70^

Studies investigating endocervical epithelial-specific changes relevant to fertility are lacking, in part due to the challenge of capturing the midcycle period when cervical mucus is permissive to sperm.^49^ Furthermore, comparisons of follicular and luteal samples based on menstrual dating or endometrial pathology can encompass substantial heterogenicity within these groupings. Meanwhile bulk transcriptome studies of the whole-organ cervix are dominated by non-epithelial cell types.^11^ Our *in vitro* study allows us to focus on epithelial changes and evaluate the effects of E2 and P4 with greater standardization, providing a cleaner evaluation of epithelial cell sensitivity to these hormones.

We used non-human primates (NHPs) due to their unique anatomical and reproductive similarities to humans. Only humans and NHPs have menstrual cycles, and lower-order animal models lack a glandular, mucus producing endocervix. However, our sample size was limited by resources and ethical considerations, and with endocervical cells harvested from four NHPs, our small sample size may have contributed to lower sensitivity in detecting significant DEGs. While we have demonstrated hormonal control of CFTR expression in prior studies,^71^ our CFTR changes here did not meet statistical significance even though expression changes trended in the same direction. There was a much closer association with E2 and P4 treated cells (Figure 4) than those in hormone-free culture, and as a hypothesis generating study, we analyzed a larger “hit” panel, recognizing that there will also be more false positives. We chose a relatively high dose of sustained E2 and a shorter P4 schedule to simulate midcycle and early luteal phase changes, respectively, and recognize that longer P4 effects may have resulted in more differentially expressed genes. Finally, *in vitro* conditions cannot fully capture physiological changes *in vivo*, including stromal-epithelial interactions,^72^ and the effects of other circulating hormones. *In vivo* transcriptomes studies are ultimately needed to further validate these findings.

## CONCLUSION

Our RNA-sequencing analysis offers a genetic overview of hormonally regulated genes related to mucus production, modification, and function within the endocervix of NHPs. This study supports findings in prior studies regarding the role of mucin-ion channel interactions, mucin post-translation modification and enzymatic activity in mucus, as well as identifies new gene targets for further functional exploration in the cervix.

